# Mechanistic insights into Zika virus NS3 helicase inhibition by Epigallocatechin-3-gallate

**DOI:** 10.1101/530600

**Authors:** Deepak Kumar, Nitin Sharma, Murali Aarthy, Sanjeev Kumar Singh, Rajanish Giri

## Abstract

Since 2007, repeated outbreaks of Zika virus (ZIKV) has affected millions of people worldwide and created global health concern with major complications like microcephaly and Guillain Barre’s syndrome. Generally, ZIKV transmits through mosquitoes (Aedes aegypti) like other flaviviruses, but reports show blood transfusion and sexual mode of ZIKV transmission which further makes the situation alarming. Till date, there is not a single Zika specific licensed drug or vaccine present in the market. However, in recent months, several antiviral molecules have been screened against viral and host proteins. Among those, (−)-Epigallocatechin-3-gallate (EGCG), a green tea polyphenol has shown great virucidal potential against flaviviruses including ZIKV. However, the mechanistic understanding of EGCG targeting viral proteins is not yet entirely deciphered except little is known about its interaction with viral envelope protein and viral protease. Since literature has shown significant inhibitory interactions of EGCG against various kinases and bacterial DNA gyrases; we designed our study to find inhibitory actions of EGCG against ZIKV NS3 helicase. NS3 helicase is playing a significant role in viral replication by unwinding RNA after hydrolyzing NTP. We employed molecular docking and simulation approach and found significant interactions at ATPase site and also at RNA binding site. Further, the enzymatic assay has shown significant inhibition of NTPase activity with an IC50 value of 295.7 nM and Ki of 0.387 ± 0.034 µM. Our study suggests the possibility that EGCG could be considered as prime backbone molecule for further broad-spectrum and multitargeted inhibitor development against ZIKV and other flaviviruses.

---

Zika virus (ZIKV), a close relative of dengue virus (DENV), is primarily a mosquito-transmitted pathogen that has already affected millions of people in more than 40 countries throughout Americas, South Pacific and South Asia(1, 2). The real danger posed by ZIKV is neurological defects like microcephaly and Guillain-Barre syndrome in newborns as well as in adults respectively(3, 4). Epidemiological studies have also reported sexual mode of ZIKV transmission is further raising the threat alarm worldwide(5). As on 1st February 2016, the World Health Organization has called a global health emergency that demands the development of safe and effective therapeutics. In 2017, WHO has confirmed three cases of ZIKV in Ahmedabad District, Gujarat, State, India (http://www.who.int/). A recent ZIKV outbreak in 2018 has been observed in India where more than 200 zika cases were confirmed including pregnant women. There is an urgency to develop antivirals against ZIKV. In past months, several bioactive molecules have been assayed either against ZIKV proteins or targeting cellular proteins by employing different approaches like screening new compound libraries or using drug repurposing(6, 7). Another essential aspect in drug discovery that could not be ignored is the use of natural products which are known to possess enormous structural and chemical variety over any other synthetic compound library(8). Moreover, natural products deliver a crucial advantage of being pre-selected evolutionary with optimized chemical structures against biological targets(9).

One of such natural products is a polyphenol called EGCG which constitutes major fraction (59 % of all polyphenols) of green tea polyphenols and has shown multiple health benefits such as antitumor, antimicrobial, antioxidative and antiviral, etc(10). The antiviral role of EGCG has been well demonstrated against several viruses such as hepatitis C virus (HCV), human immunodeficiency virus (HIV), influenza virus (FLU), DENV and chikungunya virus, etc(11–15). In a recent study, EGCG has shown a strong virucidal effect against ZIKV with a probable mechanism related to inhibiting entry into host cell demonstrated by computational finding(16, 17). However, reports suggest that apart from viral entry inhibition EGCG can also block essential steps in the replication cycle of some viruses(10). Due to the lack of complete understanding of EGCG inhibition mechanism on ZIKV, we designed our study to find a specific viral protein which could be targeted by EGCG. We have chosen NS3 helicase protein of ZIKV, a crucial enzyme in viral replication which unwinds genomic RNA after deriving energy from intrinsic nucleoside triphosphatase (NTPase) activity(18, 19). In addition to RNA unwinding activity, flavivirus helicases have also been reported to participate in other vital functions such as ribosome biogenesis, pre-mRNA splicing, RNA export and degradation, RNA maturation as well as translation etc(20). Hence, essential functions of these helicases are making them attractive drug targets.

Like other flavivirus helicases, ZIKV helicase also belongs to SF2 (Superfamily) family and a phylogenetically close relative of Murray Valley encephalitis virus (MVEV), DENV4, and DENV2(18). Full-length NS3 protein has N-terminal protease activity, and C-terminal is associated with helicase activity. ZIKV NS3 helicase (172-617 residues) is a large protein containing three domains where domain 1 (residues175-332) and domain 2 (residues 333-481) forms NTPase pocket and domain 3 (residues 481-617) in association with domain 1 and 2 forms RNA binding tunnel(21). Though ZIKV helicase is well structured, active sites at NTPase and RNA binding pockets contain highly flexible or disordered P-loop (193-203 residues) and RNA binding loop (244-255 residues) respectively, which are critical for their specific function(21, 22). In general, past decade has evidenced the significance contribution of intrinsically disordered proteins/regions (IDRs/IDPs) in almost all biological processes and the regions are considered as novel therapeutic targets (23–29). Despite the conversed active site amino acid residues among flavivirus helicases, ZIKV helicase show different motor domain movements and RNA binding modes when compared to DENV helicase(21). These vital functions of ZIKV helicase encourage to screen for antiviral molecules against its active sites. In a recent study, we have determine the inhibitory potential of a small molecule (HCQ) against ZIKV protease with computational and enzyme kinetics studies(30). Therefore, in this article, we have used molecular docking and simulation approach to find out a significant binding cavity for EGCG. Further, we have verified our computational findings by in vitro enzyme assay to probe the significant binding of EGCG at NTPase site of ZIKV helicase.

## Results

### In silico docking studies

Since for the first time, a flavivirus helicase was co-crystallized with bound ATP at substrate binding site; therefore this structure seems more significant for inhibitor screening purpose (as shown in figure 1A). Similarly, another crystal structure has ssRNA bound at helicase active site which appears suitable for employing virtual screening protocol (as shown in figure 2A). We have used extra precision (XP) mode in glide suite of Schrödinger to dock EGCG firstly at ATPase site and after that at helicase site (as shown in figure 1 and figure 2 respectively). After docking, the extent of EGCG binding at ATPase site was represented in terms of docking score as shown in table 1. A significant docking score (−7.8 Kcal mol^-1^) was observed which is contributed by various hydrogen bonding interactions with key residues of ATPase site such as ARG (202), THR (201), GLY (197), ASN (463), and ASN (417) (as shown in figure 1B, 1C and table1). Another important interaction was observed with ARG (462) which shows salt bridge and Pi-cation bonding with EGCG (figure 1B and 1C). More importantly, these interactions were reported at the critical P-loop (residues193-203) and motif VI (residues Q455, R459, and R462) of NTPase binding pocket. Mechanistic studies have already shown that P-loop residues play the most significant contribution in NTP binding and further hydrolysis(21, 31).

**Table 1:** Glide (XP) score and binding energy calculations for EGCG at ATPase and RNA binding sites.

| Active sites | Compound | Docking score (Kcal mol<sup>-1</sup>) | Binding energy (Kcal mol<sup>-1</sup>) | Ligand interactions |
| --- | --- | --- | --- | --- |
| **ATPase site** | EGCG | -7.830 | -47.324 | H-bond:ASN417, ASN463, ARG202,THR201, GLY197; Salt-bridge: ARG462; Pi-cation: ARG462 |
| **RNA binding Site** | EGCG | -7.762 | -51.312 | H-bond: ASP410, LEU430, MET414, THR225; Pi-Pi stacking: PHE289; Pi-cation: LYS431 |

**Figure 1:**
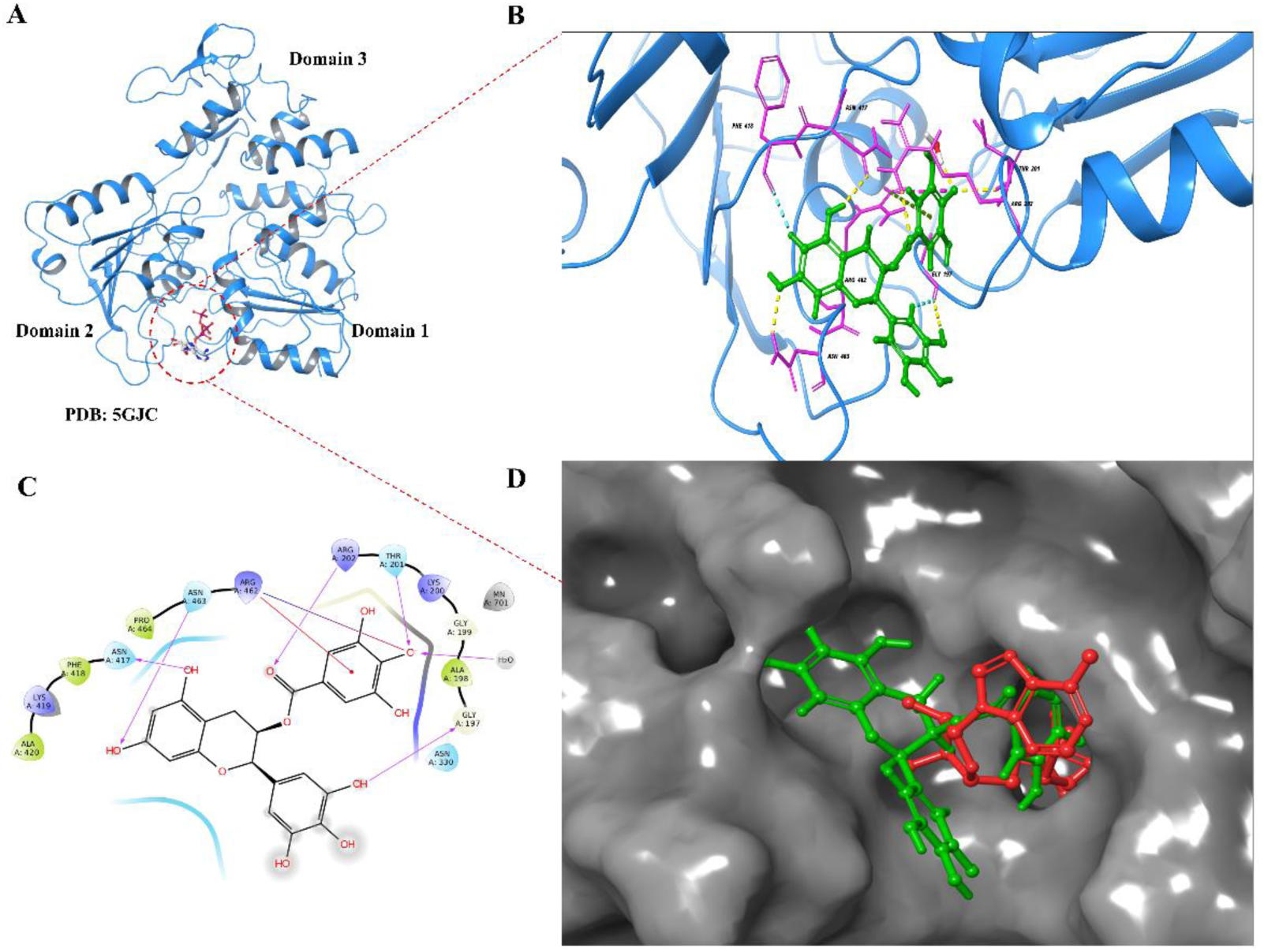
Extra precision (XP) docking of EGCG at ATPase site of ZIKV NS3 helicase. **A)** ZIKV NS3 helicase with PDB ID: 5GJC is showing ATP molecule bound at the NTPase site (red dotted circle) between domain 1 and domain 2. **B)** After molecular docking, EGCG showing molecular interactions (3D view) by H-bonds (yellow dotted lines), Pi-cation interactions (green dotted lines) and salt bridges (pink dotted lines). **C)** EGCG binding interactions were illustrated in 2D interaction diagram where interactions were represented as H-bonds (pink arrow), Pi-cation interactions (solid red line) and salt bridges (blue-red straight line). **D)** NS3 helicase represented with the solid grey surface where docked EGCG pose (green colour) was superimposed with ATP molecule (red colour) in the ATPase pocket.

**Figure 2:**
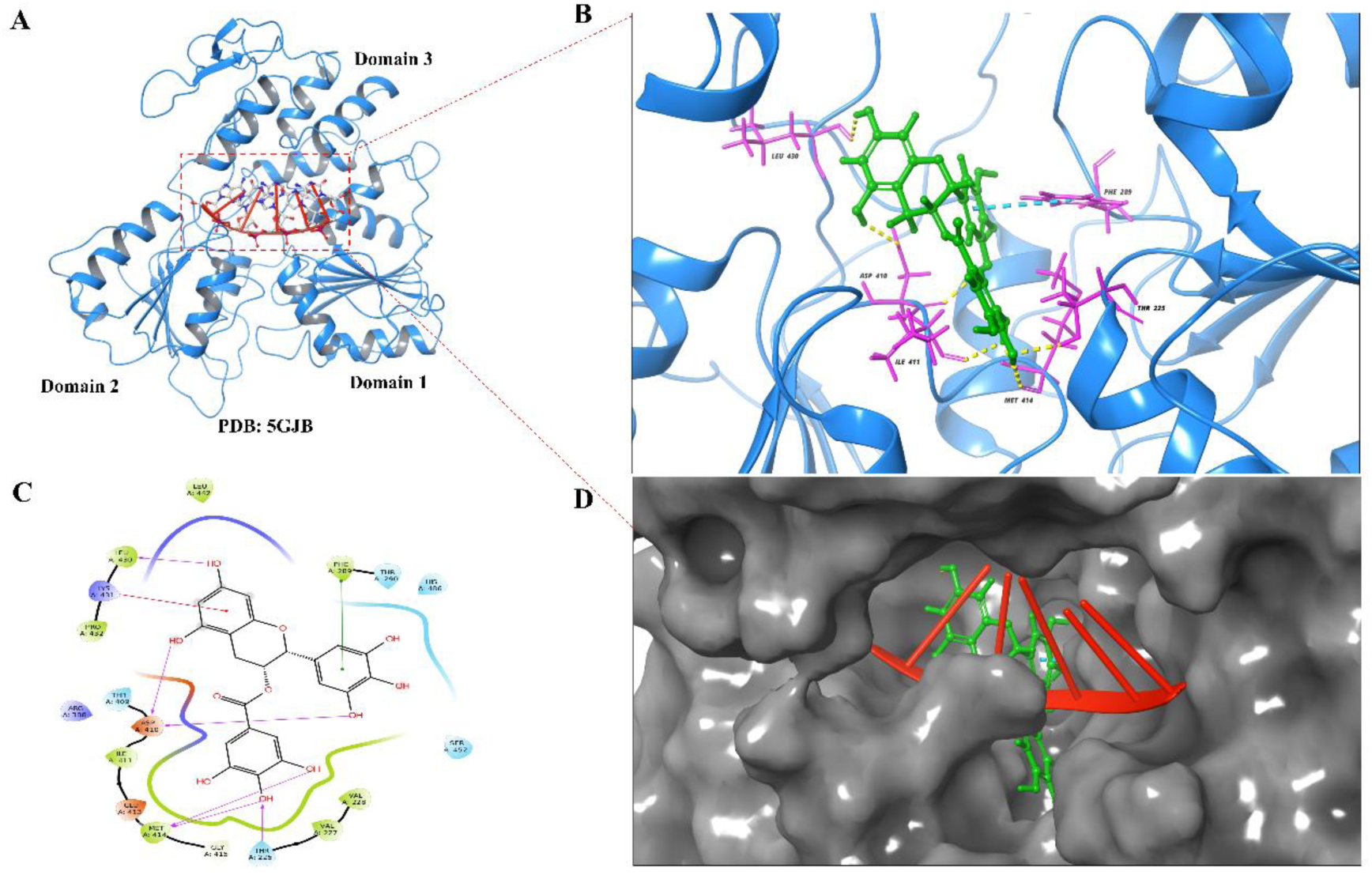
EGCG molecular docking interactions at RNA binding cavity of ZIKV NS3 helicase. **A)** NS3 helicase of ZIKV with PDB ID: 5GJB displays RNA (red colour) bound at the interface between domain 1, domain 2 and domain 3 (red dotted square). **B)** A 3D view of EGCG showing molecular interactions at the RNA binding cavity by H-bonds (yellow dotted lines), Pi-Pi interactions (Cyan dotted lines) and Pi-cation interactions (green dotted lines). **C)** 2D interaction diagram of EGCG is showing significant interaction displayed as H-bonds (pink arrow), Pi-cation interactions (red solid line) and Pi-Pi stacking (green solid lines). **D)** A solid grey surface represented by NS3 helicase and EGCG(green colour) docked pose was superimposed with RNA (red colour) bound at helicase site.

In a further study, EGCG was docked at RNA binding cavity (Figure 2) of NS3 helicase. Interestingly, EGCG was showing almost similar docking score (−7.762) as observed against ATPase site (table 1). In Figure 2B and 2C, EGCG was found to interact with RNA binding site where residues ASP 410, MET 414, LEU 430 and THR 225 were showing H-bonding with EGCG and residues LYS 431, PHE 289 were found to show Pi-cation and Pi-Pi stacking interactions respectively. From figure 2D, it can be interpreted that EGCG is binding at the entry site of the RNA molecule. Thus, it could probably interfere with helicase activity also. The residues involved in interaction with EGCG, have been shown to play an essential role in binding with RNA and further complete helicase function(21). From our docking studies, it is clear that EGCG has the potential to bind at both sites on NS3 helicase with significant interactions.

### Binding energy calculation and ADME properties

Binding energy calculations for ligand binding at protein active sites were estimated by a molecular mechanics-based approach (MM-GBSA) which employs the forcefield methods to analyze the difference in free energies of ligand, protein and the complex. The glide XP docking poses of EGCG and helicase protein at both binding sites (NTPase and RNA binding site) were used for estimating binding energies by using the Prime suite. In table1, binding energies for EGCG at RNA binding site (−51.312 kcal/mol) was shown which seems slightly higher than at ATPase site (−47.324 kcal/mol). These results show that EGCG molecule can bind at both active sites of NS3 helicase. Further, ADME properties for EGCG molecules were calculated as reported previously by Sharma et al., (2017)(17). Except, low oral absorption value, rest of the ADME properties of EGCG were within range to declare this molecule as a safe drug candidate. In support of this, a study on cell lines has reported that EGCG is only cytotoxic at concentrations greater than 200uM(16).

### Molecular dynamics simulation

#### EGCG complex at NTPase site

Molecular dynamics simulation studies help in understanding the protein structure-function such as folding, conformational flexibility, and stability. Hence, we have performed the MD simulations on the apo ZIKV helicase and compared the protein stability when EGCG bound at NTPase site and RNA binding site for a period of 100ns (Figure 3 and Figure 4 respectively). In figure 3A, the analysis of C-α root mean square deviations (RMSD) portrays that the apo ZIKV helicase represented in black did not seem to be stable enough (RMSD =1.75 – 2.75 Å) throughout the simulation when compared to the EGCG-NTPase complex represented in red (RMSD = 1.5 Å to 2.25 Å). Initially the EGCG complex was showing similar fluctuations like apo helicase but after 65 ns the complex was observed to achieve its stability till the completion of simulation course period. In order to understand the conformational fluctuations upon EGCG binding at NTPase pocket, C-α root mean square fluctuation (RMSF) at single residue level were compared between apo helicase and complex (Figure 3B). In Figure 3B, it was observed that the region around 248-255 was showng higher fluctuations of 4.75 Å in EGCG complex and lower fluctuations of 2.75 Å were monitored in region 320-326 in EGCG complex as compared to apo helicase. Notably, studies have shown that the region 248-255 belongs to RNA binding loop (244-255) which is highly dynamic and stabilized after RNA binding(21, 32). Interestingly, it was noticed that P-loop region (193-203) was not showing fluctuations in EGCG complex and apo helicase as well. Since dynamic P-loop is critical for ATP hydrolysis, it may be concluded that EGCG stabilizes the P-loop dynamics by forming significant interactions with key residues as observed in simulation interaction diagram (Figure S1C). In Figure 3C, compactness of the protein structure was measured by comparing the radius of gyration (R_g_) in apo helicase and EGCG complex. It was observed that the compactness of protein was maintained stably throughout 100ns simulation period in EGCG complex as compared to apo helicase. This also shows that the dynamic NTPase site is also controlling the overall shape of protein and EGCG is probably stabilising the NTPase site that further maintaining the overall compactness. Further, the analysis of secondary structure elements (Figure S1A) revealed that the region 180-185, 300-310, 330-350 and 450-470 shows unstable conformation throughout the simulation time. More deeply, in figure S1B the timeline of percentage index of total contacts was represented and analysis revealed that ARG 462, THR 201, GLU 286 and ARG 459 had retained the contacts throughout simulation period. In figure 3D, the interaction histogram in combination with 2D simulation interaction diagram (Figure S1D) was showing fraction of different interactions (H-bond, hydrophobic, ionic and water bridges) between EGCG and NTPase site residues. It was observed that residues GLU 286 and THR 201 have ionic interaction contribution of 100 %, while ARG 462, ARG 459 and GLY 199 shows the hydrogen bond interaction of more than 70% of simulation time. Figure 1C was showing maintenance of H-bonding throughout the simulation.

**Figure 3:**
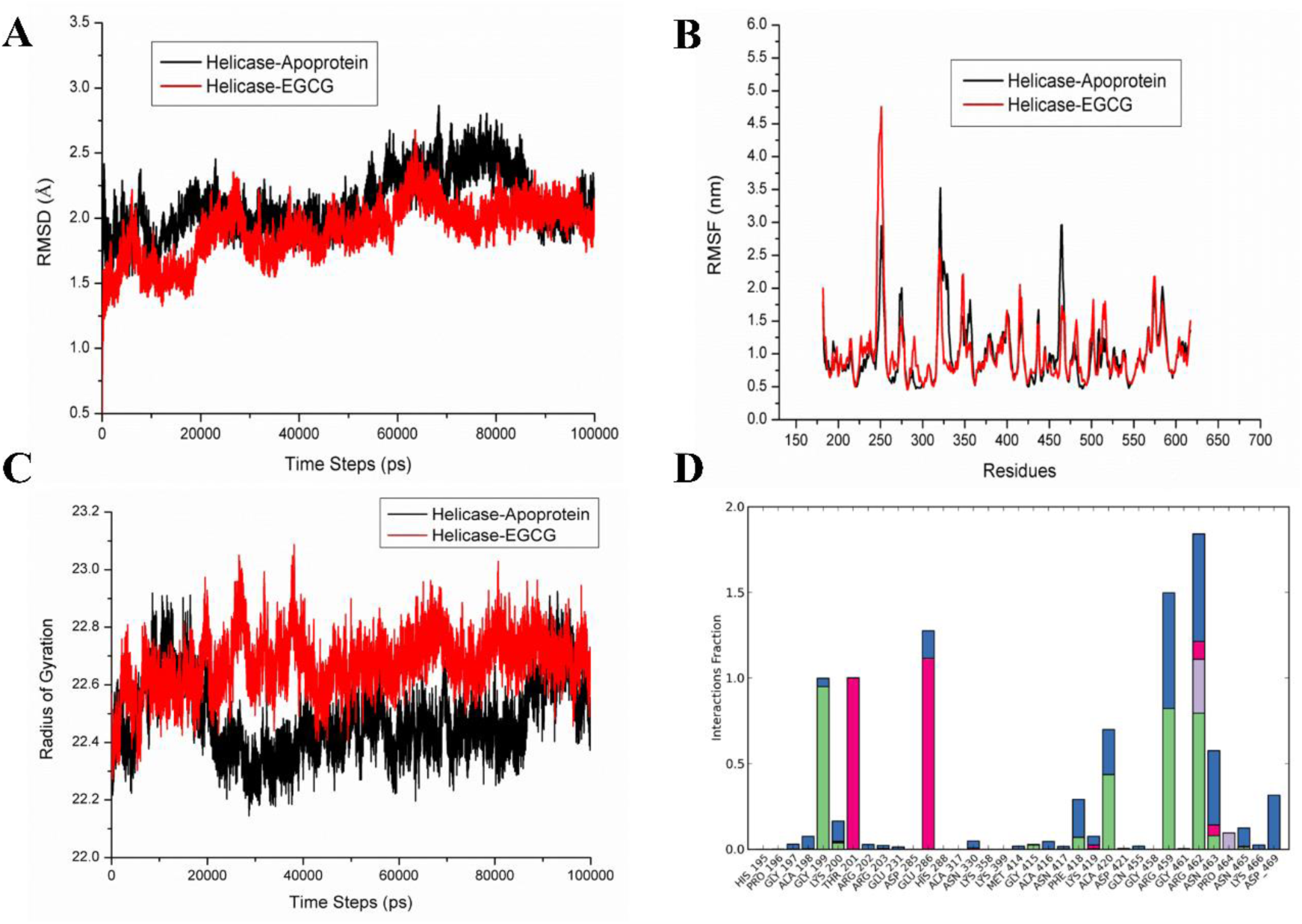
Molecular dynamics simulation of EGCG complex with ATPase site of ZIKV helicase. **A)** RMSD graph of the apo protein helicase and helicase complex with EGCG at NTPase site for the time period of 100ns simulation. **B)** The comparison of RMSF graph of Cα of the Apo protein helicase and helicase complex with EGCG at NTPase site for the time period of 100ns simulation. **C)** A comparison plot of radius of gyration of apo-NS3 helicase and EGCG complex with helicase **D)** Histogram displaying different types of interaction fractions between EGCG and ATPase site of helicase during simulation period.

**Figure 4:**
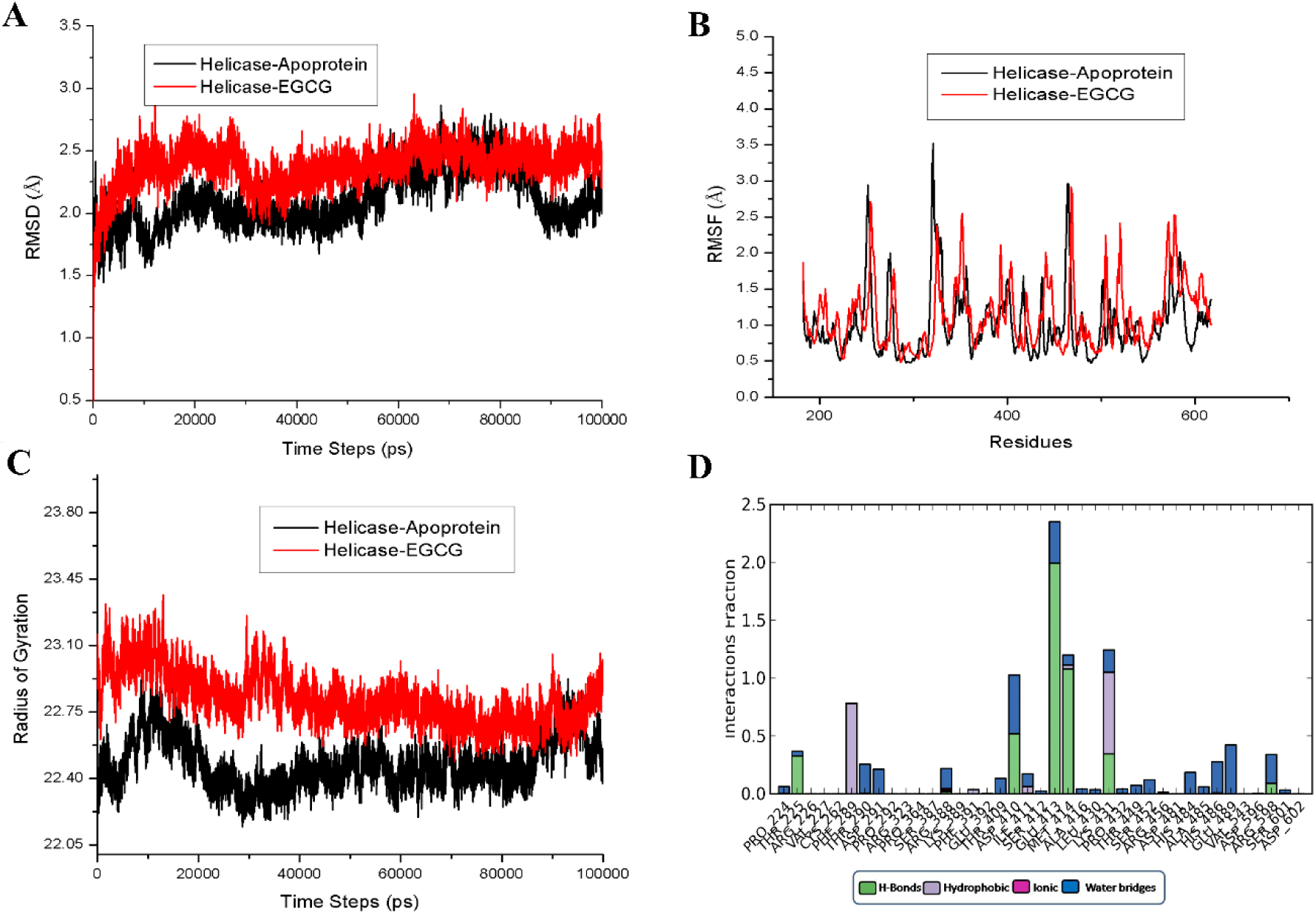
Molecular dynamics simulation of EGCG complex with RNA binding site of ZIKV helicase. **A)** RMSD graph of the apo protein helicase and helicase complex with EGCG at RNA binding site for the time period of 100ns simulation. **B)** The comparison of RMSF graph of Cα of the Apo protein helicase and helicase complex with EGCG at RNA binding site for the time period of 100ns simulation. **C)** A comparison plot of radius of gyration of apo-NS3 helicase and EGCG complex with helicase. **D)** Histogram displaying different types of interaction fractions between EGCG and RNA binding site of helicase during simulation period.

Interestingly, the metal ion Mn^2+^ which is crucial in NTP hydrolysis showed significant interaction with negatively charged oxygen atom of the EGCG (Figure S1C). In the meanwhile, the Mn2+ ion also interacts with the GLU 286 and THR 201 residue of the protein. The role of metal ion Mn^2+^ has already been established in NTP hydrolysis cycle which helps to stabilize the NTP during pre-hydrolysis step(31). Overall EGCG is showing significant interactions with crucial residues of P-loop (THR 201, GLY 199) and also with motif VI (residues ARG 459, and ARG 462) of NTPase binding pocket.

#### EGCG complex at RNA binding site

In our docking studies, we have observed that EGCG can also bind to the RNA binding cavity near entry site with significant interactions (Figure 2 and Table1). Therefore, we have run the MD simulations for apo helicase and EGCG complex at RNA binding site to compare the overall protein stability and residue level interactions (Figure 4 and Figure S2). The RMSD graph (Figure 4A) revealed that the apoprotein was showing deviation upto 20ns and attained a stable trajectory after 20ns till it reached 50ns whereas after 55ns again the deviation gradually increased by RMSD of 2.5 Å. In case of EGCG complex at RNA binding site, the deviation of complex was started increasing upto 2.5 Å till 25ns and thereafter a decrease in RMSD (2.2 Å) was observed and maintained throughout100ns period. This graph was showing that EGCG binding at the RNA site stabilising the overall protein conformation. In figure 4B, the comparison of RMSF was revealing that the higher fluctuations of almost 3.5 Å were observed in region 320-326 in apo helicase which were further reduced to 2.0 Å in EGCG-RNA site complex. Interestingly it was noticed that region 248-255 was showing higher fluctuations of 4.75 Å in EGCG complex at NTPase site (Figure 3B) in comparison to EGCG complex at RNA site (Figure 4B) which was showing similar fluctuations of 2.75 Å like apo helicase. Overall, the RMSF plot of EGCG-RNA site complex was found to support RMSD graph. As already mentioned before, region 248-255 contains crucial RNA binding loop which is important for helicase activity(21). From above comparisons, it may be interpreted as EGCG binding at NTPase site may lead to the conformational changes at RNA binding loop region (244-255) which otherwise were not observed when EGCG bound at RNA site. Further in figure 4C, the radius of the gyration plot was showing that EGCG complex at RNA site had maintained compactness throughout the 100ns period as compared to apo helicase. In figure 4D, the different types of interaction fractions were observed that showed mostly H-bonded interactions were prominent between EGCG and protein. In Figure S2A, the extent of formation of secondary structure elements was analysed when EGCG bound to RNA site throughout 100ns simulation period. Further, it was noticed that H-bonding was maintained throughout the simulation (Figure S2C) and mostly residues GLU 413, MET 414 were continuously contacted throughout while residues LYS 431, PHE289 and ASP 410 were showing irregular contacts (Figure S2B and S2D). This residues observed in interaction with EGCG at RNA site, have essential role already mentioned in crystal structure of helicase with RNA(21). For example, LYS410 and ASP410 shows important interaction with RNA sugar bases. Hence our simulation study shows that EGCG may have the capability to significantly bind at RNA site also along with NTPase site.

### *In-vitro* experiments

#### Inhibition of NTPase activity

NTPase activity inhibition by EGCG has already been reported in literature against bacterial DNA gyrases(33). Our molecular docking and simulation studies have shown that EGCG can bind to both the active sites (NTPase and RNA binding site) of ZIKV NS3 helicase with significant interactions between critical residues. Since NTP hydrolysis provides the required energy to open up RNA secondary structures during replication, we firstly focused on the NTPase activity inhibition assays experimentally.

The *E. coli* expressed recombinant NS3 helicase (53.6kDa) was purified by Ni-NTA affinity chromatography as shown in figure 5A. ATPase activity of NS3 helicase was determined by absorbance (630nm) based malachite green method which estimates the release of free phosphate. The Michaelis-Menten equation was used to quantitate the kinetic parameters (K_m_, K_cat_ and K_cat_/K_m_) for NTPase activity of NS3 helicase. The K_m_, K_cat_ and K_cat_/K_m_ were calculated as 345.9 ± 30.31 µM, 68.96 ± 1.71 min^-1^ and 0.1993 ± 0.056 µM^-1^ min^-1^ respectively (Figure 6B). Inhibition assays were carried out in triplicates and enzyme is preincubated for 10 minutes with varying concentrations of EGCG. Further, we have observed the dose dependent inhibition of NTPase activity and IC50 values were calculated as 295nM against NTPase site (Figure 5C). Additionally, the inhibitory potential of EGCG was determined using substrate velocity curves which showed inhibitory constant (K_i_) of 0.387 ± 0.034 µM (Figure 5D). This result shows that EGCG molecule is quite capable of inhibiting NTPase activity of NS3 helicase in low micromolar range and could act as potential lead backbone molecule for further inhibitor development.

**Figure 5:**
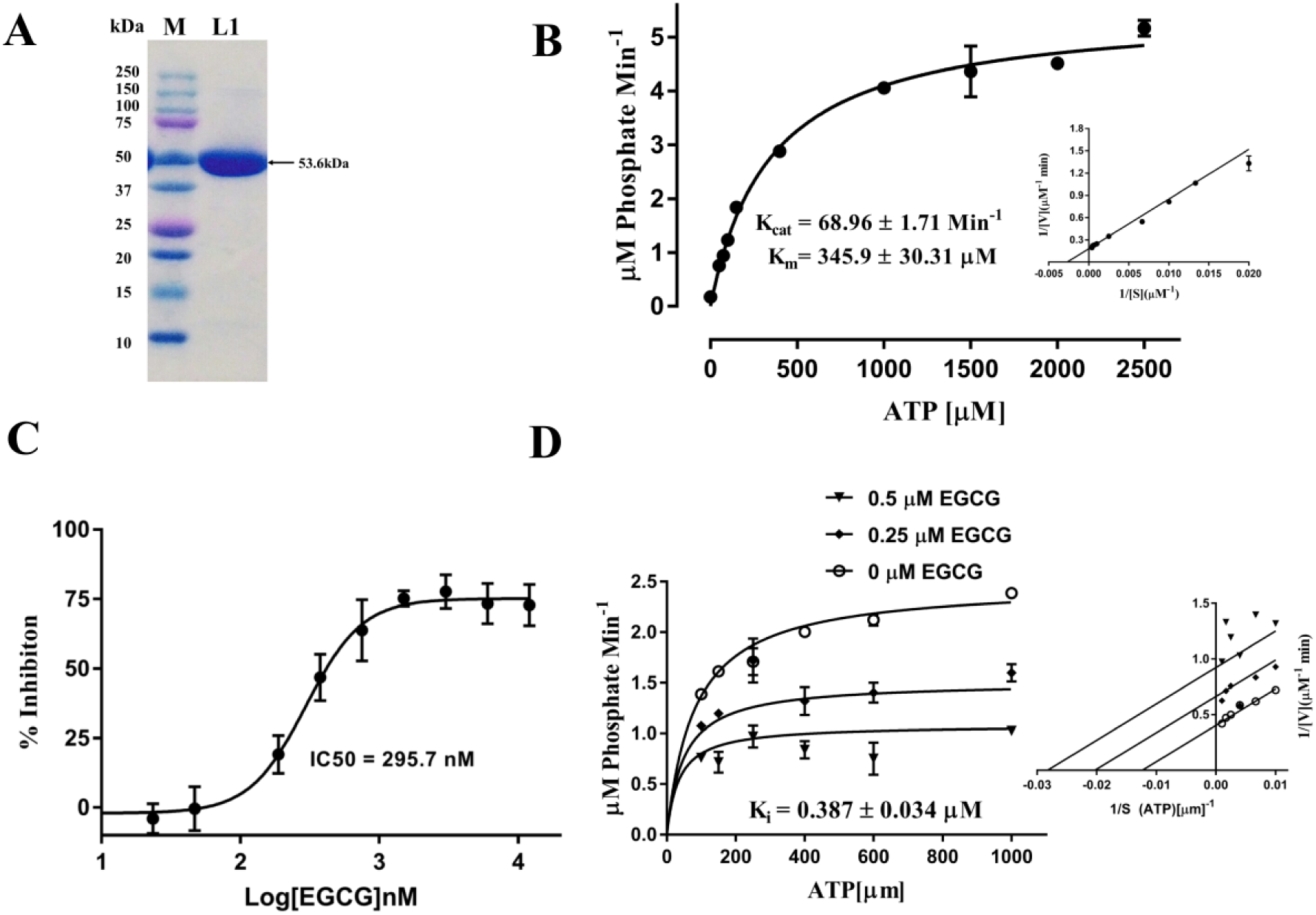
EGCG inhibits ATPase activity of NS3 helicase. **A)** Purification of NS3 helicase by Ni-NTA affinity chromatography. A 10% SDS gel was run and stained with Coomassie dye where final his-tagged NS3 helicase fractions were pooled and concentrated by Amicon (10kDa) centrifugal filter. In this figure, M-protein ladder (Bio-Rad Precision Plus) and L1 contains purified concentrated his-tagged NS3 helicase protein of ZIKV **B)** Kinetic parameters (Substrate-Velocity curve) calculated for 80 nM NS3 helicase after varying the substrate (ATP) concentration ranging from 50 µM to 2500 µM. **C)** IC50 calculated for EGCG against NTPase site by incubating 80 nM helicase with varying concentration of EGCG (serially diluted: 1200 nM: 23.40 nM) **D)** Inhibition constant (Ki) was calculated for 80nM helicase at different concentration of ATP (100, 150, 250, 400, 600 and 1000 µM) and EGCG concentrations were kept at 0.5 and 0.25 µM.

## Discussion

In recent years, repeated outbreaks of ZIKV has necessitated the urgent need for developing specific drugs. Also, the major complications of ZIKV infections are related to pregnant women, therefor it is important to find molecules which are safe and have minimal or no side-effects. Considering the safety point, natural products have always been a great source of drugs or drug like molecules and also these molecules have evolutionary pre-optimized biological targets(8). To find specific biological targets, in silico structure-based drug discovery approaches have revolutionized and fasten the current drug developing strategies. In fact, the molecules which can target specifically viral proteins could act as safe therapeutics against ZIKV(34). EGCG, a green tea polyphenol has shown significant antiviral activity against several viruses including HIV, HSV, CHIKV and some flaviviruses like HCV, and DENV(10). Recently, in ZIKV, EGCG inhibitory potential was determined in a cell line based study where probable mechanism was related to interaction of the compound with envelope protein(16). Previously, we have also supported the EGCG envelope protein interaction with computational study(17). However, reports suggested that EGCG may target other viral proteins which are important in genome replication and maturation(10). Due to lack of adequate experimental support regrading EGCG envelope protein interactions and considering the possibility of finding more specific target for EGCG, we have chosen NS3 helicase protein of ZIKV for determining potential inhibitory effects of EGCG. NS3 helicase of ZIKV is an attractive drug target due to its essential role in opening RNA secondary structures during replication(21). Also, reports suggest that EGCG has shown anti-ATPase activity against bacterial DNA gyrases(33).

It is well known fact that flavivirus helicases are motor proteins and require energy released from NTP hydrolysis to perform their helicase function(35). Therefore, firstly we analysed the EGCG affinity towards NTPase site through docking, binding energy calculation and MD simulations. These studies revealed that EGCG can dock significantly with key residues (ARG 202, THR 201, GLY 197, ASN 463 and ASN 417) at NTPase site and further MD simulations were supporting the stable EGCG interaction with residues (Mn^2+^, ARG 462, THR 201, GLU 286 and ARG 459) were carried out throughout the simulation period (100ns). In crystal structure of ZIKV NS3 helicase with bound ATP, these residues have significant functions such as: the Mn^2+^ co-ordination with GLU 286 stabilizes the ATP molecule; the P-loop residues (GLY 197, ARG 202, LYS 200) and motif VI residues (ARG 459, ARG 462) are playing key role ion NTP hydrolysis by interacting with transition state nucleotides(21).

Recently, a mutational study has shown that residues THR 201, ARG2 02, and GLU 286 are critical for NTP hydrolysis by ZIKV NS3 helicase(31). Also, a compound NITD008 was shown to inhibit ZIKV replication experimentally where mechanism was elucidated computationally to show binding at the NTPase site with significant interactions at P-loop region(36). Based upon computational findings, we have done inhibition assays where EGCG has shown significant dose dependent inhibition of NTPase activity with IC50 of 295.7 nM by Malachite green method. Further, the mode of inhibition was determined to be uncompetitive with inhibition constant (K_i_) in low micromolar range (K_i_ = 0.387 ± 0.034 µM). A study of polyphenols inhibiting ZIKV protease has shown that EGCG inhibits protease with higher IC50 (87 µM) values(37). Taken together, our findings suggest that EGCG may target ZIKV helicase more specifically in addition to envelope protein and NS3 protease.

In ZIKV NS3 helicase the RNA binding site along with NTPase site has flexible pockets containing critical loop regions needed to perform function(21, 22). Therefore, we also studied the possibility of EGCG interacting at RNA binding site. This was due to the fact that polyphenols have shown potential interactions with intrinsically disordered regions in proteins and could be seen as novel strategies of drug development against IDPs(38). Also, literature has shown that viral proteins have several short stretches of disordered regions within proteins and more propensity of intrinsically disordered active sites(22, 24, 39). Our docking and MD simulations studies are showing that EGCG has the ability to bind at the entry site of RNA binding pocket with significant interactions. More specifically, EGCG is showing different types of interactions (H-bond, ionic, salt bridge) with residues GLU 413, MET 414, LYS 431, PHE289 and ASP 410. In crystal structure, these residues are playing key role in binding to RNA(21). In our MD simulations, it has also reported that EGCG binding at NTPase site is increasing fluctuations in RNA site which could interpreted as an allosteric relationship between two sites. This observation could be supported by the recent study on DENV helicase where NTPase and RNA sites show allosteric effects.

In summary, our extensive docking and simulation analysis is showing that EGCG can bind strongly to the NTPase site and can inhibit the activity of ZIKV NS3 helicase more precisely supported by in-vitro enzyme kinetics assays. Also, EGCG can form the significant binding interactions at RNA site revealed by computational tools. Interestingly, the comparison with previous studies illustrate that EGCG can target multiple viral protein such as envelope(16), protease(37) and now more precisely helicase. Since EGCG has shown virucidal effect against several viruses, therefore EGCG backbone could be used to develop a broad-spectrum antiviral molecule in near future.

## Materials and methods

### In silico docking studies

In-silico studies were carried out on X-Ray crystal structure (PDB ID: 5GJC, resolution 2.2 Å) of NS3 helicase bound to ATP-Mn^2+^ and crystal structure containing bound ssRNA with PDB ID: 5GJB (1.7 Å). Further, the docking process was initiated by following a series of necessary steps such as protein preparation, receptor grid generation, ligand preparation and finally ligand docking. Protein preparation was carried out by using Protein Preparation wizard in Schrodinger LLC Maestro v11.0. In protein preparation, force field OPLS-2005 was utilized for H-bond network optimization and energy minimization. Co-crystallization artifacts such as missing side chains and loops were filled by using Prime. Protonation states were generated using Epik at pH 7.4. In literature, three crystal water molecules were reported at the substrate binding site, therefore except these water molecules rest were deleted beyond 5 Å from the ligand. A receptor grid was generated on the centroid of substrate binding site by picking up the atoms of co-crystallized ligand (ATP) in 5GJC and ssRNA in 5GJB. The length of the grid was kept 20 Å. Before final docking step, EGCG molecule was also prepared using LigPrep module of Maestro as described by Sharma et. al. Further, ligand docking was carried out by using extra precision glide (Glide XP) program from Schrodinger (Glide, Version 11). EGCG molecule was docked flexibly at rigid active site on the protein. Final XP dock scores were analyzed for significant interactions with active site residues.

#### Binding energy calculation and ADME properties

Prime/MM-GBSA approach was used to calculate the free energy of binding for XP docked complex. This method was utilized as described previously by Sharma et al., (2017)(17). MM/GBSA is an empirical scoring that approximates the ligand binding affinities with the receptor. Following equation is used for calculating the free energy of binding-ΔG (bind) = E_complex(minimized) - (E_ligand(minimized)+E_receptor(minimi zed) QikProp module of Schrödinger software (QikProp, version 4.3, Schrodinger) was used for the calculation of the drug like behavior through the evaluation of the pharmacokinetic properties that are required for the absorption, distribution, metabolism, and excretion (ADME)(17). These properties have been calculated already in our previous study(17).

#### Molecular dynamics simulation

Molecular dynamics simulation was executed for the apoprotein and complex of ZIKA helicase with the EGCG at NTPase site and RNA binding site using the Desmond module implemented in Schrodinger(a). The OPLS-AA (Optimized Potentials for Liquid Simulations – All Atom) 2005 force field was used for the minimization of the complex and apoprotein(40). The structures of the protein and complex were imported in the Desmond setup wizard and were solvated in a cubic periodic box of TIP3P water molecules. The structures were neutralized by adding a suitable number of counter ions and 0.15 M of salt concentration(41). Steepest descent, a hybrid method is implemented for the local energy minimization of the system. The limited memory Broyden-Fletcher-Goldfarb-Shanno algorithm with a maximum of 5000 steps is used until a gradient threshold (25 kcal/mol/Å) were reached. The constant NPT (number of atoms, Pressure P and temperature T) ensemble condition is incorporated to relax the simulation system to generate simulation data for post analyses. The overall simulation process is performed using the Nose-Hoover thermostats, and stable atmospheric pressure (1atm) carried out by Martina-Tobias-Klein barostat method, and the 300K was assigned as the temperature value. In order to investigate the equation of motion throughout the dynamics, the multi-time step RESPA integrator algorithm was used(42, 43). The bonded, near non-bonded and far non-bonded interactions, were assigned at the time steps of 2,2 and 6fs respectively. The atoms involved in the hydrogen bond interaction were constrained with the SHAKE algorithm. A cut-off value of 9 Å radius was set up to estimate the long-range electrostatic interactions and Lennard-Jones interactions. The Particle Mesh Ewald (PME) method was used to evaluate the long-range electrostatic interactions along with the simulation process using the periodic boundary conditions (PBC). During the intervals of 1.2 and 4.8ps, the trajectory data and the energy analysis were recognized. The final production molecular dynamics were carried out for 100ns for both the apoprotein and protein complex. The results were analyzed using the simulation event analysis and simulation interaction diagram available in Desmond module(44).

## In-vitro experiments

### Cloning, Expression and Purification

ZIKV NS3 helicase coding region (1342 bp) corresponding to PDB ID: 5GJC was synthesized by GeneArt Gene synthesis services provided by Invitrogen (USA). This gene was further ligated into pET 151/D-TOPO vector purchased from Thermo Fisher Scientific (USA). The final construct containing N-terminal 6X-His tag with TEV protease cleavage site was transformed into BL21 (Sigma) *E*.*coli* cells and thereafter positive clones were expressed in LB broth media (inducing with 1mM IPTG at 20°C overnight). Cells containing recombinant protein were harvested by centrifugation at 6000g at 4°C and re-suspended in binding buffer (50mM Tris, 300mM NaCl, 40mM imidazole, 5%glycerol pH8.0). Protease inhibitor cocktail (Thermo Fisher Scientific, USA) was added before cell lysis. Cells were lysed by sonication and adding 50% B-PER (Thermo scientific) reagent. After centrifugation at 16000 rpm for 30 min at 4°c, the supernatant containing recombinant protein was filtered and loaded on HisTrap FF 5ml column (GE healthcare). Recombinant protein was eluted by using a linear imidazole gradient from 0 to 100 %. Eluted fractions were analyzed on 10% SDS-PAGE for the purity. Buffer exchange was carried out to remove the Imidazole by using Amicon 10 kDa (Merck Millipore) centrifugal filters. Recombinant protein was kept in storage buffer (50mM Tris, 100mM NaCl, 5% glycerol) stored in aliquots at -80°C.

### NTPase Activity Assay

The NTPase activity of recombinant NS3 helicase was analyzed by using malachite green method as described previously in literature(45, 46). We have little modified the ratio of reagents as; 1mg/ml Malachite green, 2mg/ml Ammonium molybdate, 0.7 M HCl and 0.05 % Triton X-100. All the reagents were prepared in ultrapure water ensuring that there is no phosphate contamination. A blank sample O.D. below 0.3 at 630 nm (TECAN infiniteM200PRO) confirms the phosphate free assay system. A phosphate standard curve (serially diluted 6.25 µM - 100 µM) was prepared (40 µL sample + 160 µL Malachite reagent) which is used further to quantitate the amount of free phosphate released by NS3 helicase. In 96 well micro-plate assay, NS3 helicase was pre-incubated in duplicates at a concentration of 80 nM in 20 µl assay buffer (40 mM Tris, 80 mM NaCl, 8 mM Mg(AcO)2, 1 mM EDTA, pH 7.5). The NTP hydrolysis reaction was started after adding 10 µL ATP in varying concentrations from 25 µM to 2500 µM. Final sample volume was kept 40 µL and after 20 minutes of incubation at 25°C, reaction was terminated by adding 160µL of malachite reagent. After incubating the reaction at room temperature for 5 minutes absorbance was measured at 630 nm. All the kinetic parameters were calculated by plotting the data in GraphPad Prims software 7.0. The data fitting was done using Michaelis-Menten equation (V= V_max_[S]/ (K_m_+ [S]) to calculate kinetic parameters (V_max_, K_m_ and K_cat_).

### NTPase Activity Inhibition Assay

Inhibition assays were carried out in similar buffer conditions as used in activity assay. EGCG (Sigma-Aldrich) was dissolved in water at a stock concentration of 5mM. Inhibition assay was carried out in triplicates in 96 well plate. Initially, the 80nM enzyme is pre-incubated with a varying concentration of EGCG (serially diluted:12000 nM to 23.40 nM) in assay buffer at 25°C for 10 minutes. After incubation, 1mM ATP substrate was added to the wells and incubated for 20 minutes. Finally, the reaction was stopped by adding 160 uL malachite green reagent to all the wells. Absorbance was taken at 630 nm after 5 minutes. IC50 value was calculated by fitting the data in non-linear regression mode using GraphPad Prism 7.0. Further, the inhibition kinetic parameters were calculated at different ATP concentrations as 100, 150, 250, 400, 600 and 1000 µM. Two concentrations of EGCG were chosen in 40 µl sample volume as 0.5 µM and 0.25 µM which are above and below from IC50 value. For all the reactions, 80 nM enzyme was pre-incubated in assay buffer with different EGCG concentrations for 10 minutes. Afterward ATP was added in varying concentrations as mentioned above and incubated for 20 minutes. Malachite reagent (160 µL) was added in each well and absorbance was taken at 630 nm after 5 minutes. All the measurements were taken in triplicates and Ki (Inhibition constant) was determined after fitting data in GraphPad Prism 7.0.

## Funding & Acknowledgment

This work was partially supported by DST grant, India (YSS/2015/000613) to RG and IIT-Mandi, India to RG. DK is grateful to ICMR fellowship.

## Author’s Contributions

RG, conception, design and study supervision; DK, NS, AM, and SKS D.K.: acquisition, analysis, and interpretation of data, writing, and review of the manuscript.

