## Supplementary Files for "Mechanistic insights into Zika virus NS3 helicase inhibition by Epigallocatechin-3-gallate"

### Supplementary figures

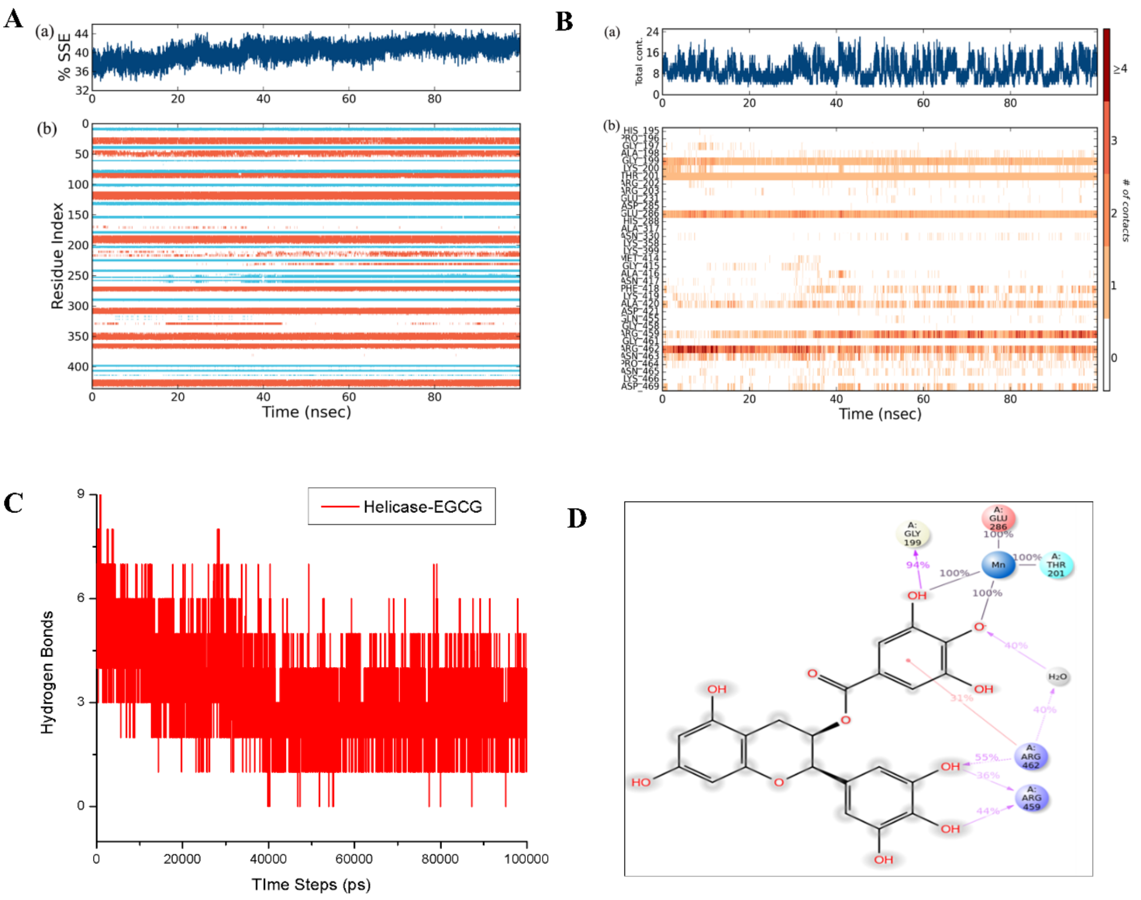


**Figure S1: MD simulation analysis for EGCG complex at ATPase site of helicase. A)** Secondary structure elements analysis of ZIKV helicase - EGCG complex at NTPase site for a time period of 100ns simulation. Blue colour representation shows beta-sheet conformation, orange colour shows the alpha-helix conformation and white colour represents the loop conformation. **B)** Percentage analysis of total contacts during the simulation period of 100 ns. **C)** Hydrogen bond analysis of the ZIKV helicase – EGCG complex at NTPase site. **D)**2-Dimensional Interaction diagram of the ZIKV Helicase – EGCG complex at NTPase site after 100ns simulation.

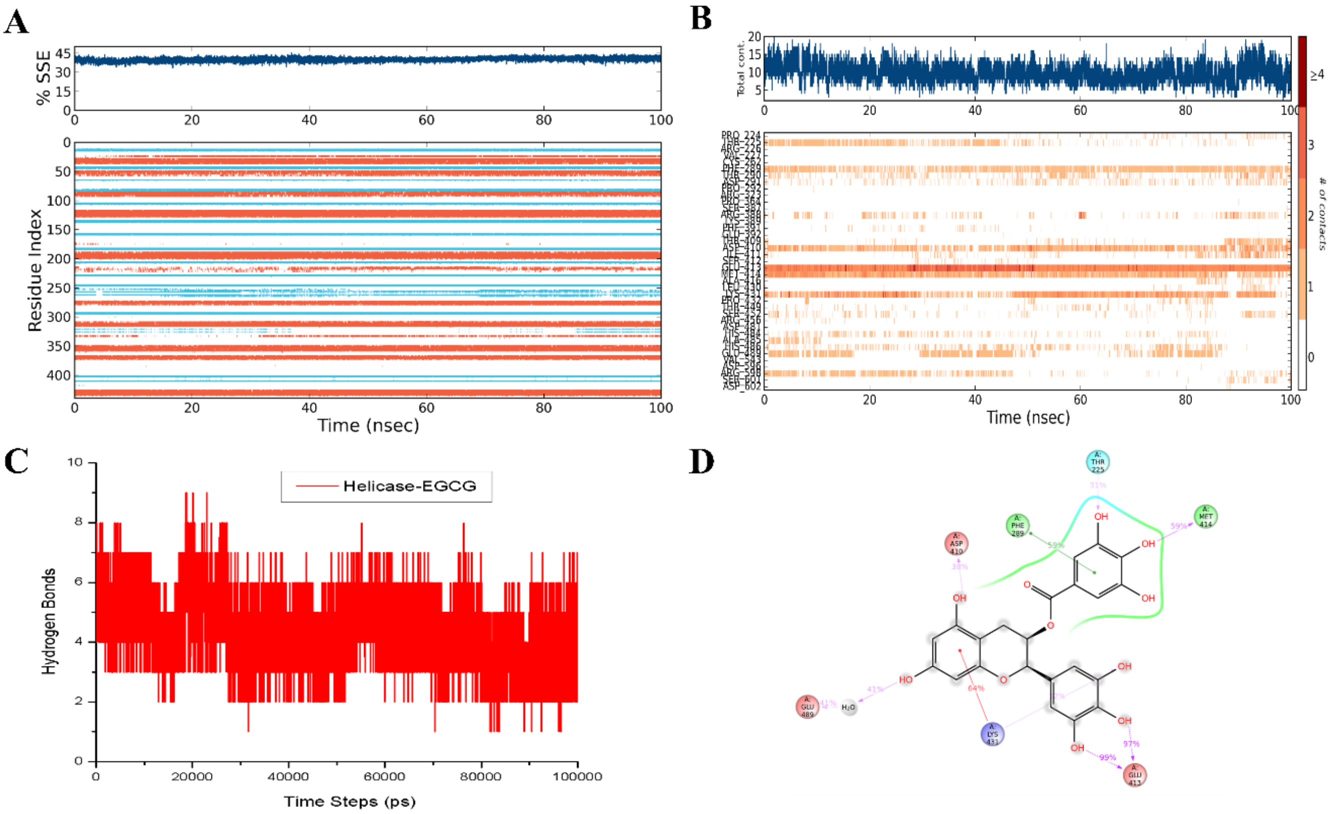


**Figure S2: MD simulation analysis for EGCG complex at RNA binding site of helicase.** **A)** Secondary structure elements analysis of ZIKV helicase - EGCG complex at RNA binding cavity for the time period of 100ns simulation. Blue colour representation shows beta-sheet conformation, orange colour shows the alpha-helix conformation and white colour represents the loop conformation **B)** Percentage analysis of total contacts during the simulation period. **C)** Hydrogen bond analysis of the ZIKV helicase – EGCG complex at RNA binding site. **D)** 2-Dimensional Interaction diagram of the ZIKV Helicase – EGCG complex at RNA binding site after 100ns simulation.
